# Pan-TREM-1 versus macrophage-restricted TREM-1 blockade in cancer and other inflammatory pathologies

**DOI:** 10.1101/2025.08.22.671865

**Authors:** Alexander B. Sigalov

## Abstract

**Background:** Triggering receptor expressed on myeloid cells 1 (TREM-1), an inflammation amplifier, is an emerging target in inflammation and oncology.

**Objective:** To test my hypothesis that pan-TREM-1 and macrophage-restricted TREM-1 blockades may differ in their efficacy in cancer and other inflammatory diseases.

**Methods:** Ligand-independent TREM-1 inhibitory peptides GF9 and GA31 (the latter in a form of macrophage-targeted lipopeptide complexes, GA31-LPC) were used as pan-TREM-1 and macrophage-restricted TREM-1 inhibitors, respectively, to test the hypothesis in multiple animal models of cancer, sepsis, pulmonary inflammation and fibrosis.

**Results:** In fully immunocompetent mice, GA31-LPC but not GF9 overcomes pancreatic cancer resistance to PD-L1 blockade and synergizes with immunotherapy. In PANC-1 xenograft-bearing athymic nude mice, GF9 and GA31-LPC both increase complete response rate and survival when combined with chemotherapy but exhibit opposing dependence on timing of treatment initiation. GF9 is effective only when given with but not after chemotherapy. In contrast, GA31-LPC is effective only when given after but not together with chemotherapy. Critical dependence of the therapeutic efficacy of TREM-1 blockade on the type of blocker and treatment timing was also observed in animal models of sepsis and acute lung injury but not fibrosis.

**Conclusion:** This study provides the first evidence that pan-TREM-1 and macrophage-restricted TREM-1 blockades may strikingly differ in their therapeutic efficacy depending on the disease, timing of treatment initiation and the type and stage of inflammatory response. This opens new avenues for development of TREM-1-targeting strategies and leads to a new framework for the treatment of cancer and other inflammation-associated diseases.

## 1. Introduction

First reported in 2000 (1), triggering receptor expressed on myeloid cells 1 (TREM-1) was initially shown to play a role in sepsis (2). Currently, TREM-1 is well recognized as a key player in cancer (3–4) and numerous other inflammation-associated diseases and disorders of infectious and non-infectious origin (reviewed in (5–9). Upon inflammation, TREM-1 is upregulated and amplifies inflammatory response by mediating release of proinflammatory cytokines and factors (8, 10–11), functioning as a switch between physiological and pathophysiological inflammatory processes.

In animal models, TREM-1 blockade ameliorates cancer (reviewed in (3, 12–13), sepsis (reviewed in (14–15), acute respiratory distress syndrome (ARDS) and other acute lung injuries (16–18), inflammatory bowel disease (11, 19–20), retinopathy (21), gouty arthritis (22), rheumatoid arthritis (RA) (23–25), atherosclerosis (26–27), empyema (28), brain and spinal cord injuries (29–34), Parkinson’s disease (35), liver diseases (36–40), renal injury and kidney transplantation (41–42), pertussis (43), skin fibrosis (44), pulmonary fibrosis (PF) (45), hemorrhagic shock (46), reperfusion injury (47), and thrombosis (48), implicating TREM-1 as a potential "magic bullet" in the treatment of diseases with underlying inflammatory pathologies.

TREM-1 is mainly expressed on neutrophils, monocytes, and macrophages including monocyte-derived macrophages (1, 49). Different types of cells that express TREM-1 can play different or even opposite roles in the pathogenesis of inflammatory diseases (50–54) depending on the disease and the type and stage of inflammatory response. This led me to hypothesize that in these cases the therapeutic activity and efficacy of pan-TREM-1 inhibitors that target TREM-1 on all TREM-1-expressing cells can strongly differ from those of cell-restricted TREM-1 inhibitors that target TREM-1 on a certain type of cells that express TREM-1 (e.g., macrophages). Despite more than two decades of intensive research in the field, this question has never been addressed before mostly, due to the lack of cell-restricted approaches to TREM-1 blockade.

Current TREM-1 inhibitors can be broadly classified into ligand-dependent and ligand-independent based on their mechanism of action (**Figure 1A**). Ligand-dependent inhibitors all attempt to block interactions of TREM-1 with its multiple known and unknown ligands by binding either to TREM-1 ligands (e.g., decoy peptides LP17 and LR12) or to the receptor itself (e.g., an anti-TREM-1 monoclonal antibodies, mABs; human eCIRP-derived peptide M3; PGLYRP1 protein-derived peptide N1 and small molecule VJDT) (**Figure 1A**) (57). In terms of cell specificity, ligand-dependent inhibitors (**Figure 1A**) are all "pan-TREM-1" inhibitors since they inhibit TREM-1 on all cells that express TREM-1 (**Figure 1B**).

**Figure 1.**
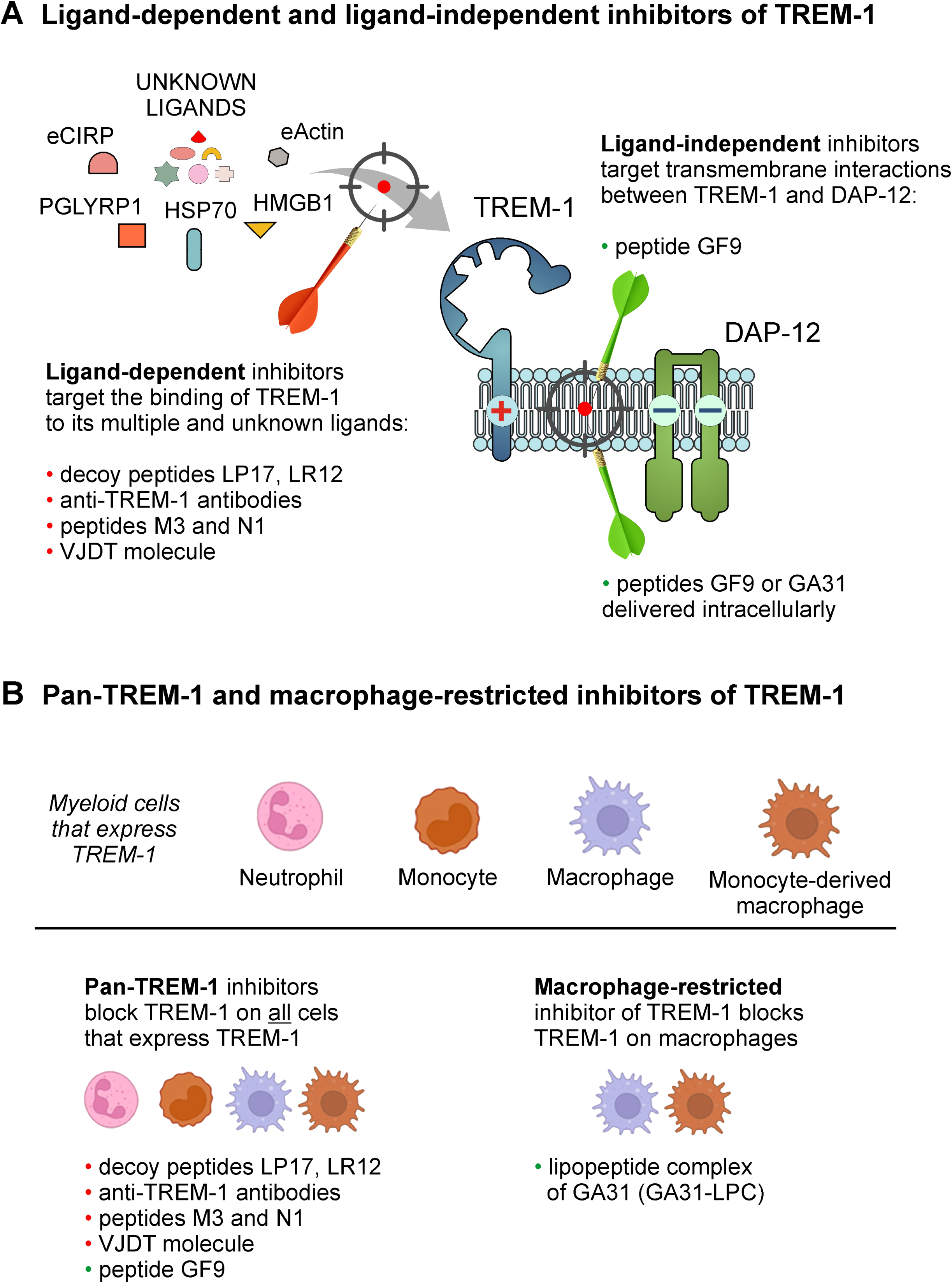
Classification of TREM-1 inhibitors based on their mechanisms of action and cell specificity. **(A)** Currently available TREM-1 inhibitors can be classified into two groups according to their mechanisms of action: ligand-dependent and ligand-independent inhibitors. TREM-1/DAP-12 receptor complex assembly is depicted where TREM-1 and DAP-12 are bound in the cell membrane by electrostatic interactions. Ligand-dependent inhibitors attempt to block interaction of TREM-1 with its multiple and unknown ligands (shown by red arrow) while ligand-independent inhibitors disrupt the interactions between TREM-1 and DAP-12 in the cell membrane (shown by green arrow) that blocks transmembrane signaling. **(B)** Based on their cell specificity, TREM-1 inhibitors can be classified as pan-TREM-1 inhibitors that inhibit TREM-1 on all cells that express TREM-1 and cell-restricted inhibitors that inhibit TREM-1 on a certain type of TREM-1-expressing cells (i.e., macrophages). Abbreviations: PGLYRP1, peptidoglycan recognition protein 1; eCIRP, extracellular cold-inducible RNA-binding protein; HMGB1, high mobility group box 1; Hsp70, heat shock protein 70 kDa.

Ligand-independent approach to TREM-1 blockade is principally different and uses structural features of TREM-1. TREM-1 has no intracellular signaling domain and signals via its partner, DAP12 (1, 58) (**Figure 1A**). As such, TREM-1 belongs to the family of the so-called multichain immune recognition receptors (MIRRs) with ligand-recognizing and signal-transducing domains located on separate subunits (chains) (59). First reported in 2004 (59–60), Signaling Chain HOmoOLigomerization (SCHOOL)-driven molecular mechanisms of MIRR signaling revealed the transmembrane protein-protein interactions between MIRR ligand-recognizing and signal-transducing subunits as therapeutic targets that can be targeted by short synthetic peptides (SCHOOL peptides) (59, 61–62). When applied to TREM-1, this led to discovery of the TREM-1 inhibitory SCHOOL peptide sequence GF9 that inhibits TREM-1 by disrupting the interactions between TREM-1 and DAP-12 in the membrane (**Figure 1A**). This inhibitory action has been shown to ameliorate cancer and many other inflammatory diseases and conditions *in vivo* (25, 44, 62–66).

Molecular mechanisms of ligand-independent TREM-1 blockade allow to develop not only pan-TREM-1 but also cell-restricted TREM-1 inhibitors. When systemically delivered as a free peptide, GF9 reaches its site of action in the cell membrane from outside the cell (**Figure 1A**). GF9 lacks cell specificity and inhibits TREM-1 on all cells that express TREM-1 and as such represents a pan-TREM-1 inhibitor (**Figure 1B**). To develop a macrophage-restricted TREM-1 inhibitor, the peptide GA31 containing the peptide sequence GF9 at the N-terminal end has been designed and incorporated into macrophage-specific nanosized lipopeptide complexes, GA31-LPC (25, 62). When intracellularly delivered by GA31-LPC to macrophages via scavenger receptor A (SR-A)-mediated endocytosis (36), GA31 is released and reaches its site of action in the cell membrane from inside the cell (**Figure 1A**). Thus, by inhibiting TREM-1 predominantly on macrophages, GA31-LPC represents a macrophage-restricted TREM-1 inhibitor (**Figure 1B**).

Here, GF9 and GA31-LPC were used to comparatively study pan-TREM-1 and macrophage-restricted TREM-1 blockades in animal models of cancer, sepsis, pulmonary inflammation and fibrosis. I show for the first time that the therapeutic activity and efficacy of TREM-1 blockade critically depend on the type of TREM-1 inhibitor used, the disease, the animal model studied, timing of treatment initiation and the type and stage of inflammatory response. This finding opens new avenues toward our understanding of the TREM-1-mediated inflammatory response in human health and disease and leads to a new framework for the treatment of cancer and other inflammation-related diseases. A lack of consideration of this previously unrecognized phenomenon may cause serious misinterpretations of data of preclinical and clinical studies evaluating the efficacy of inflammation-targeting strategies including discovery and development of therapeutic modulators of TREM-1 signaling.

## 2. Materials and methods

### 2.1. TREM-1 inhibitory peptides and formulations

Previously reported TREM-1 inhibitory peptides GF9 and GA31 (25, 44, 62) were synthesized by AmbioPharm, Inc. (North Augusta, SC, USA). Macrophage-specific LPC formulations containing GA31 were generated essentially as previously reported (25).

### 2.2. *In vivo* mouse studies

Animal experiments were performed by various contract research organizations (CROs), including Translational Drug Development (TD2; Scottsdale, AZ, USA), Washington Biotechnology, Inc. (WBI; Baltimore, MD, USA), Crown Bioscience (San Diego, CA, USA),, Noble Life Sciences (Sykesville, MD, USA), Charles River Laboratories (Wilmington, MA, USA), and the Center for Translational Medicine at Thomas Jefferson University (Philadelphia, PA, USA) on a fee-for-service basis.

#### 2.2.1. Cancer

In xenografts studies in female athymic nude mice (Crl:NU(NCr)-Foxn1^nu^), the mice were inoculated subcutaneously in the right flank with 0.1 mL of a 50% DMEM / 50% Phenol Red-free Matrigel mixture containing a suspension of 5 x 10^6^ cells/mouse of PANC-1 tumor cells. Twenty-six days following inoculation, mice with tumor volumes of 71-159 mm^3^ were randomized into groups of ten mice, each with a group mean tumor volume of 107-108 mm^3^ by random equilibration. PANC-1 xenograft-bearing mice were treated intraperitoneally (i.p.) with vehicle (phosphate-buffered saline, PBS, pH 7.4), 25 mg/kg GF9 or 13 mg/kg GA31-LPC in combination with standard-of-care (SOC) chemotherapy (80 mg/kg gemcitabine i.p. Q3Dx4 and 30 mg/kg nanoparticle albumin based paclitaxel (nab-ptx) intravenously, i.v., Q3Dx4) starting either Day 1 or Day 13 and then continuing as post-chemo therapy (QDx7 every other week to the end of study at Day 99). Tumor volumes and body weights were recorded when the mice were randomized and two times weekly thereafter. The relative tumor volume (RTV) was calculated using the following formula: RTV = (tumor volume on measured day)/(tumor volume on day 0). All results were expressed as the median values (n = 10). Complete response was identified as tumor volume of less than 13.5 mm^3^ for three consecutive measurements.

In syngeneic orthotopic studies in female C57BL/6 wild-type (WT) mice, the mice were inoculated into the subcapsular region of the pancreas with a mPA6115-Luc [luciferase-expressing Kras (G12D)/Trp53 null/Pdx1-cre (KPC)] tumor chunk. Mice were IVIS imaged for bioluminescence on Day 3 and assigned to the respective treatment groups based on the bioluminescent photon flux. Mice were dosed within 24 hours of randomization. Treatment with vehicle (PBS, pH 7.4), GF9 (25 mg/kg, once a day for 3 weeks, i.p.) or GA31-LPC (13 mg/kg, once a day for 3 weeks, i.p.) alone and in combination with immunotherapy, an anti-mouse PD-L1 antibody (10 mg/kg, twice a week for 3 weeks, i.p.), was initiated after grouping and continued throughout the experiment. Tumors were measured by bioluminescent imaging twice a week starting Day 4. The study was terminated on Day 23.

#### 2.2.2. Sepsis

In endotoxic shock studies in male C57BL/6 WT mice, the mice challenged with lipopolysaccharide (LPS) essentially as described previously (67) were treated i.p. once with vehicle (PBS, pH 7.4), 25 mg/kg GF9 or 13 mg/kg GA31-LPC 1 h prior or 1 or 3 h post LPS. In positive control group, mice were treated i.p. once with 0.1 mg/kg dexamethasone 1 h prior LPS.

In cecal slurry injection (CSI) studies in female Balb/C mice, the mice challenged with CSI essentially as described (68) were treated i.p. once with vehicle (PBS, pH 7.4), 25 mg/kg GF9 or 13 mg/kg GA31-LPC 1 h prior or 6 or 12 h post CSI challenge. In positive control group, mice were treated i.p. with 25 mg/kg ceftriaxone and 12.5 mg/kg metronidazole (ABX) preventively 1 h prior CSI, then daily for 5 days.

#### 2.2.3 Pulmonary inflammation and fibrosis

In acute lung injury studies in male Sprague Dawley (SD) rats, the rats were treated with LPS from E.coli O26:B6 dosed via oropharyngeal aspiration (o.p.) once on Day 1 (0 hr). In the prevention model, rats were dosed i.p. once with vehicle (PBS, pH 7.4), 25 mg/kg GF9 or 13 mg/kg GA31-LPC 1 h prior LPS. In the treatment model, rats were dosed i.p. once with 25 mg/kg GF9 or 13 mg/kg GA31-LPC 1 h post LPS challenge. All animals were euthanized 6 h post LPS challenge. Lungs were lavaged with sterile PBS (without calcium and magnesium) to collect bronchoalveolar lavage fluid (BALF). BALF (per animal) were weighed and total and differential BALF cell counts were determined in cell pellets.

In PF studies in C57BL/6 WT mice, the mice were treated with bleomycin (BLM) intratracheally (i.t.) once on Day 0. In the prevention model, mice were dosed i.p. daily with vehicle (PBS, pH 7.4), 25 mg/kg GF9 or 13 mg/kg GA31-LPC for two weeks starting Day 1. At Day 7, half of mice from each group were sacrificed. In the treatment model, mice were dosed i.p. daily with vehicle (PBS, pH 7.4) 25 mg/kg GF9 or 13 mg/kg GA31-LPC for two weeks starting Day 15. After euthanization (Days 7 and 22 in the prevention model or Day 28 in the treatment model), the left lungs were lavaged with sterile PBS to collect BALF. The right lungs were harvested and analyzed for lung hydroxyproline.

### 2.3. Statistical analysis

For comparative analyses between two groups of data, statistically significant differences were assessed by Student’s unpaired t-test for normally distributed variables. When the assumption of Gaussian distribution was not met, a non-parametric Mann-Whitney U-test was used for comparisons. For comparisons between more than two groups, statistical differences were analyzed with the one- or two-way analysis of variance (ANOVA) test. A p-value < 0.05 was considered statistically significant: *p < 0.05; ** p < 0.01; *** p < 0.001 and **** p < 0.0001. Statistical calculations and graphs were done using GraphPad Prism software.

## 3. Results

### 3.1. Anti-tumor efficacy of GF9 and GA31-LPC oppositely depends on timing of treatment initiation relative to chemotherapy in cancer mice with intact innate immunity but lacking T cells

Similar to other cancer treatments (e.g., surgery, radiation, radiopharmaceuticals, antibody-drug conjugates, etc.) (69–72), current SOC chemotherapies, including gemcitabine/nab-ptx and FOLFIRINOX (71), represent a double-edged sword: they reduce tumor burden by killing cancer cells, however, the resulting dead tumor cells, or debris, induce inflammation that may lead to the failure of therapy, cancer recurrence and metastasis (69, 72–74). This suggests that timely resolution of therapy-induced inflammation may improve response to treatment as well as prevent tumor recurrence and metastasis resulting in improving quality of life and overall survival of patients.

In this study, to demonstrate that dampening of the TREM-1-mediated inflammation induced by cancer treatment can suppress cancer progression and improve response rate, the anti-tumor efficacy of GF9 and GA31-LPC and its dependence on treatment timing relative to chemotherapy were evaluated in subcutaneous PANC-1 xenograft-bearing athymic nude mice. Selection of this model was based on two considerations: (a) TREM-1 is involved in innate immunity (6), and (b) athymic nude mice retain a fully functional innate immune system (natural killer cells, macrophages, neutrophils) but lack mature T cells (75), which makes the model especially suitable for studying modulators of innate immunity such as TREM-1 inhibitors.

In combination with chemotherapy, GF9 and GA31-LPC both synergistically inhibit tumor growth, improve survival (not shown) and up to three times increase the complete response rate as compared to chemotherapy alone but with opposite dependence of the efficacy outcomes on the timing of treatment initiation relative to chemotherapy (**Figure 2A**). While GF9 is effective when given with but not after chemotherapy, GA31-LPC is effective only when given after chemotherapy but not with chemotherapy. As in our previous studies in mice bearing other pancreatic cancer xenografts (76), ligand-independent TREM-1 blockade by using GF9 and GA31-LPC was well tolerable in this study (data not shown).

**Figure 2.**
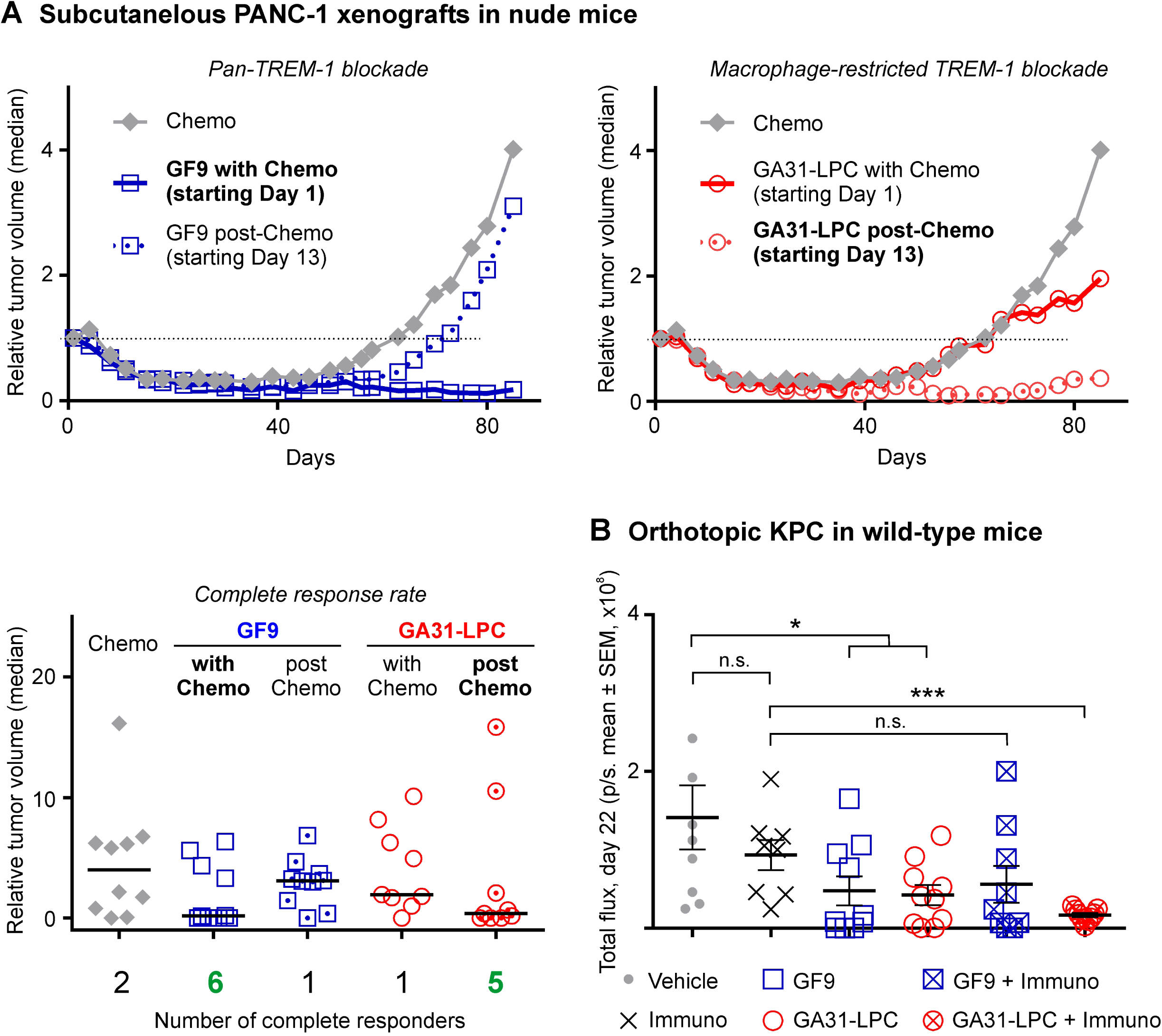
Comparative anti-tumor efficacy of GF9 and GA31-LPC alone and in combination with chemotherapy or immune checkpoint blockade. **(A)** In athymic nude mice, a pan-TREM-1 inhibitor GF9 significantly inhibited PANC-1 tumor growth and improved the complete response rate when given with but not post-chemotherapy. In contrast, a macrophage-restricted TREM-1 inhibitor GA31-LPC was effective when given post-chemotherapy but not with it. The relative tumor volume (RTV) was calculated using the following formula: RTV = (tumor volume on measured day)/(tumor volume on day 0). The study was terminated on Day 99. All results are expressed as the median values (n = 10). **(B)** In fully immunocompetent mice, GA31-LPC and GF9 alone both inhibited orthotopic KPC tumor growth. In combination with PD-L1 blockade, GA31-LPC but not GF9 overcame tumor resistance and synergized with immunotherapy. Tumors were measured by IVIS bioluminescent imaging twice a week starting Day 4. The study was terminated on Day 23. Complete response was identified as tumor volume of less than 13.5 mm^3^ for three consecutive measurements. Abbreviation: KPC, luciferase-expressing Kras (G12D)/Trp53 null/Pdx1-cre tumor. *, p<0.05; ***, p<0.01; n.s., not significant.

In summary, these findings indicate that combination of chemotherapy with pan-TREM-1 blockade or macrophage-restricted TREM-1 blockade yielded a significant synergistic anti-tumor effect compared to monotherapy. However, the effect was critically and oppositely dependent on the timing of GF9 or GA31-LPC treatment initiation relative to chemotherapy, thereby providing the first experimental *in vivo* evidence that the anti-tumor effect of TREM-1-mediated modulation of the innate immune response caused by cancer therapy strongly depends on the cell specificity of the modulator used and treatment timing.

### 3.2. GA31-LPC but not GF9 overcomes tumor resistance to PD-L1 blockade and synergizes with immunotherapy in fully immunocompetent cancer mice

To determine whether pan-TREM-1 and macrophage-restricted TREM-1 blockade alone or in combination with immune checkpoint blockade (ICB) are similarly effective against cancer in fully immunocompetent mice, C57BL/6 WT mice with orthotopically inoculated KPC tumor were treated with GF9 or GA31-LPC alone or in combination with anti-PD-L1 antibody.

In this study, GF9 and GA31-LPC alone both significantly inhibited KPC tumor progression (**Figure 2B**). These findings are in line with the literature data which revealed that GF9 alone inhibits tumor growth and improves survival in C57BL/6J WT mice with orthotopically implanted hepatocellular carcinoma (HCC) Hep55.1C (37) and Hepa1.6 (66) tumors.

In agreement with well-known limitations of anti-PD-L1 ICB for the treatment of solid tumors in general (77–78) and the ineffectiveness of immunotherapy in pancreatic cancer specifically (79), in this study, PD-L1 blockade alone did not affect KPC tumor growth (**Figure 2B**). Treatment of cancer mice with GF9 concurrently with immunotherapy did not exhibit any synergistic anti-tumor activity compared to GF9 alone (**Figure 2B**). In contrast, the combination of PD-L1 blockade with GA31-LPC synergistically suppressed tumor growth compared to GA31-LPC alone (**Figure 2B**).

Thus, these data demonstrate that while TREM-1 blockade alone either with GF9 or GA31-LPC is effective in suppressing tumor progression, only macrophage-restricted TREM-1 inhibitor GA31-LPC, but not pan-TREM-1 inhibitor GF9 could overcome tumor resistance to PD-L1 blockade and synergize with immunotherapy.

### 3.3. GA31-LPC but not GF9 effectively protects septic mice from death with opposing dependence on timing of treatment initiation relative to challenge

Therapeutic effect of TREM-1 blockade in experimental sepsis was first demonstrated in 2001 (2). Currently, TREM-1 is widely recognized as a critical contributor to the immune dysfunction caused by sepsis (14–15).

To explore whether GF9 and GA31-LPC differ in their ability to protect septic animals against death and whether the protection level depends on the mode of TREM-1 blockade (prevention or treatment), two models were used in this study: mice with endotoxic shock induced by LPS and mice with polymicrobial sepsis induced by CSI.

In endotoxemic mice, a slight but still not significant (p=0.1) effect on the survival rate was observed in animals dosed once with GF9 preventively (1 h prior LPS), while therapeutic administration of GF9 (1 or 3 h post LPS) seemingly did not affect animal survival (**Figure 3A**). These findings are in line with the previously observed decrease of effectiveness of TREM-1 blockade in experimental sepsis at later times of treatment with the highest level of protection in the prevention models (2). In contrast to GF9, GA31-LPC significantly protected the animals from death when administered once either preventively 1 h prior LPS or therapeutically 1 h or 3 h post LPS (**Figure 3A**). The exhibited efficacy of GA31-LPC surprisingly did not decline at later times of treatment (1 or 3 h post-LPS challenge) but rose paradoxically compared to that of GA31-LPC administered preventively (1 h prior LPS) (**Figure 3A**). Further, no significant differences in the survival rate were noted between mice treated therapeutically with GA31-LPC and mice treated preventively with dexamethasone (positive control) (**Figure 3A**).

**Figure 3.**
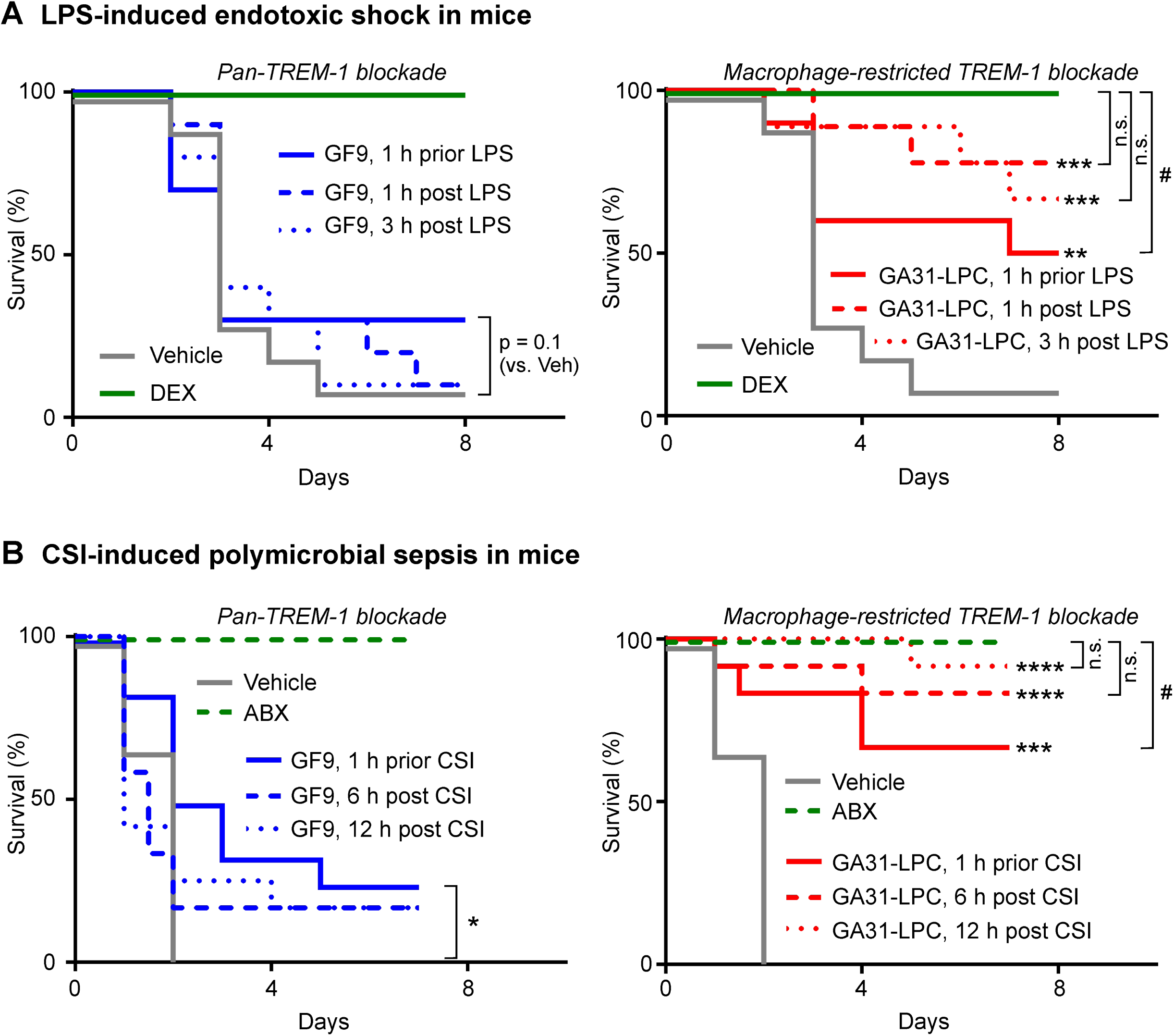
Comparative efficacy of GF9 and GA31-LPC in protecting mice from death caused by endotoxic shock or polymicrobial sepsis. **(A)** In lipopolysaccharide (LPS)-challenged C57BL/6 wild-type (WT) mice, a pan-TREM-1 inhibitor GF9 given once 1 h prior LPS showed a slight but still not significant effect while no effect was observed when GF9 was given once 1 or 3 h post-LPS. In contrast, a macrophage-restricted TREM-1 inhibitor GA31-LPC significantly protected the animals from death when administered once preventively (1 h prior LPS) and therapeutically (1 or 3 h post LPS). The exhibited efficacy of GA31-LPC given at 1 or 3 h post-LPS did not differ from that shown by positive control (dexamethasone, DEX, 1 h prior LPS) but was paradoxically higher compared to that of GA31-LPC given preventively. **, p<0.01 (versus vehicle); ***, p<0.001 (versus vehicle); ^#^, p<0.05 (versus DEX); n.s., not significant. **(B)** In cecal slurry injection (CSI)-challenged C57BL/6 WT mice, GF9 given once preventively (1 h prior CSI) but not therapeutically (6 or 12 h post-CSI) showed a slight significant effect. In contrast, GA31-LPC significantly protected the animals from death when administered once preventively (1 h prior CSI) and therapeutically at 6 or 12 h post CSI. The exhibited efficacy of GA31-LPC given at 6 or 12 h post-CSI challenge did not differ from that shown by positive control (ceftriaxone and metronidazole, ABX, beginning 1 h prior CSI, then daily for 5 days) but was paradoxically higher compared to that of GA31-LPC given preventively. *, p<0.05 (versus vehicle); ***, p<0.001 (versus vehicle); ****, p<0.0001 (versus vehicle); ^#^, p<0.05 (versus ABX); n.s., not significant.

In line with the data in LPS-challenged endotoxic mice, in mice with polymicrobial sepsis induced by CSI, a slight but significant effect on the survival rate was observed in animals dosed once with GF9 preventively (1 h prior CSI) but not therapeutically (6 or 12 h post CSI) (**Figure 3B**). In contrast, GA31-LPC significantly protected the animals from death when administered once either 1 h prior or 6 or 12 h post CSI (**Figure 3B**). Similar to the data in endotoxemic mice, the exhibited efficacy of GA31-LPC did not decline at later times of treatment (6 or 12 h post-CSI) but rose compared to that of GA31-LPC administered preventively (1 h prior CSI) (**Figure 3B**). Further, no significant differences in the survival rate were noted between mice treated therapeutically with GA31-LPC and mice treated with ceftriaxone and metronidazole (positive control) beginning 1 h prior CSI, then daily for 5 days.

In summary, these data highlight a marked contrast between pan-TREM-1 and macrophage-restricted TREM-1 blockades in their ability to protect endotoxic or septic mice from death.

Notably, they also reveal a paradoxical enhancement in the therapeutic activity of GA31-LPC treatment when administered as a treatment rather than as a preventative measure. These findings strongly support further development of this promising therapeutic approach to significantly improve survival of septic patients.

### 3.4. GA31-LPC but not GF9 significantly suppresses acute pulmonary neutrophilia in rats

Neutrophil infiltration into the lung is a hallmark of acute lung injury (ALI) and its most severe manifestation, ARDS (80). In animal models of LPS-induced ALI, TREM-1 blockade ablates neutrophilic lung inflammation (16–17).

Since TREM-1 blockade efficacy in experimental sepsis significantly and differently depended on the type of TREM-1 inhibitor (GF9 versus GA31-LPC) and timing of treatment initiation relative to challenge (**Figure 3**), I hypothesized that this may also be the case in experimental ALI. Indeed, in this study, the data in rats with LPS-induced pulmonary neutrophilia strongly support the hypothesis. GF9 did not affect BALF neutrophil content when given either preventively (1 h prior LPS) or therapeutically (1 h post LPS) (**Figure 4A**). In contrast, GA31-LPC significantly decreased neutrophil infiltration into the lungs when given 1 h post LPS but was ineffective when given 1 h prior LPS (**Figure 4A**). While surprising, this finding is consistent with the data obtained in experimental sepsis showing that preventive treatment with GA31-LPC is not as effective as therapeutic intervention (**Figure 3**).

**Figure 4.**
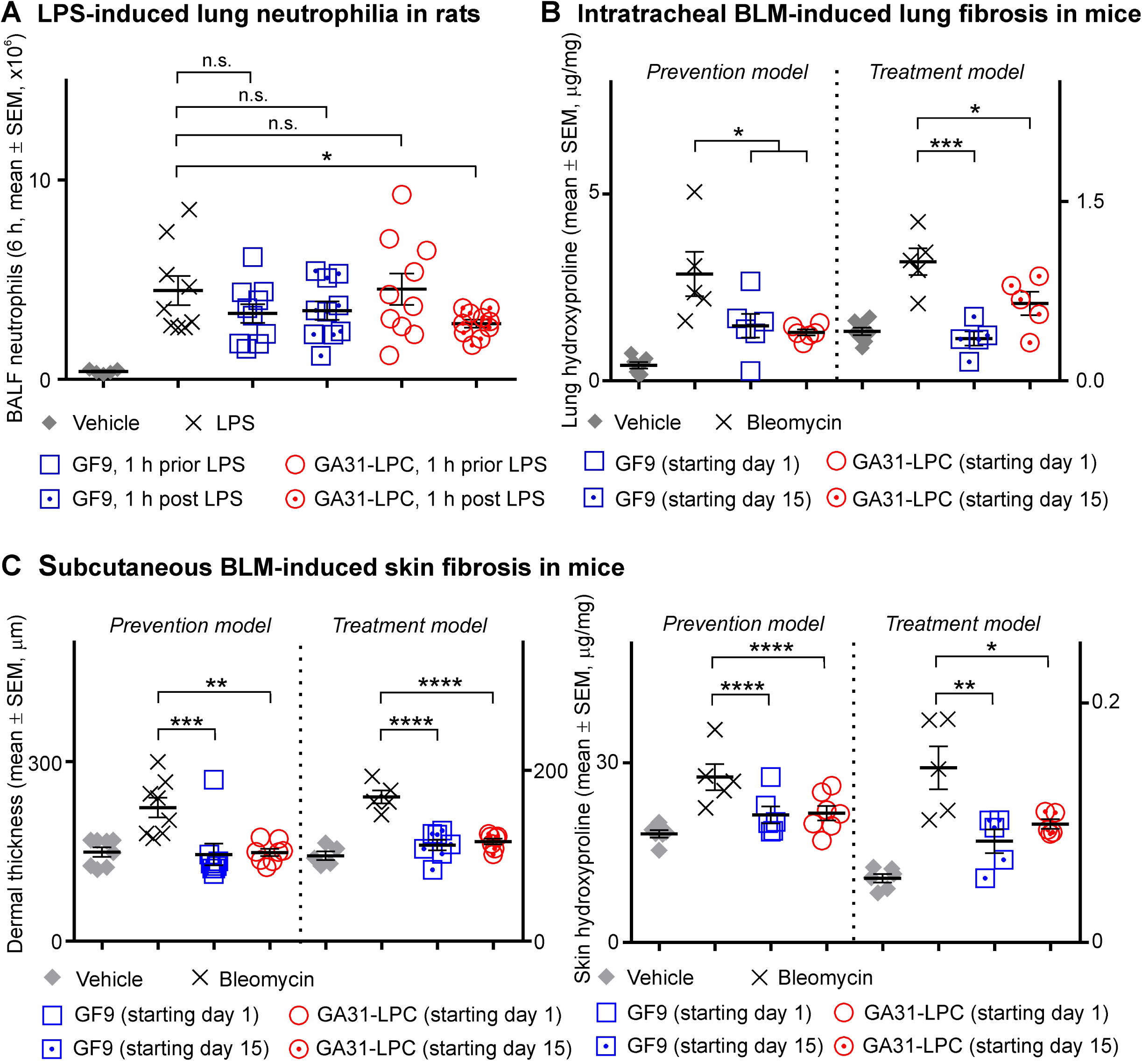
Comparative efficacy of GF9 and GA31-LPC in preventing and treating lipopolysaccharide-induced acute lung injury in rats as well as bleomycin-induced pulmonary and skin fibrosis in mice. **(A)** In Sprague Dawley (SD) rats with acute lung neutrophilia induced by oropharyngeally aspirated lipopolysaccharide (LPS), a pan-TREM-1 inhibitor GF9 did not affect bronchoalveolar lavage fluid (BALF) neutrophil content when given either preventively (1 h prior LPS) or therapeutically (1 h post LPS). In contrast, a macrophage-restricted TREM-1 inhibitor GA31-LPC significantly decreased neutrophil infiltration into the lungs when given as a therapeutic (1 h post LPS), but surprisingly not as a preventive (1 h prior LPS) agent. **(B)** In C57BL/6 wild-type (WT) mice with intratracheal bleomycin (BLM)-induced pulmonary fibrosis, GF9 and GA31-LPC were similarly effective in preventing and reversing fibrosis as analyzed by lung hydroxyproline. **(C)** In C57BL/6 WT mice with subcutaneous BLM-induced skin fibrosis, GF9 and GA31-LPC were similarly effective in preventing and reversing fibrosis as analyzed by dermal thickness (left panel) and skin hydroxyproline (right panel). Adapted from Bale S, Verma P, Yalavarthi B, Bajzelj M, Hasan SA, Silverman JN, et al. JCI Insight (2024) 9:e176319. *, p<0.05; **, p<0.01; ***, p<0.001; ****, p<0.0001; n.s., not significant.

Thus, these findings mirror the trend observed in our animal studies of cancer, endotoxemia and sepsis, which implies that pan-TREM-1 and macrophage-restricted TREM-1 blockades may strongly differ not only in their ability to prevent and/or treat acute inflammatory conditions, but also in how their therapeutic efficacy depends on the timing of treatment initiation.

### 3.5. GF9 and GA31-LPC both effectively prevent and reverse pulmonary fibrosis in mice

TREM-1 plays a role in PF (81) and is overexpressed in the lungs of mice with BLM-induced PF (82). In this study, pan-TREM-1 and macrophage-restricted TREM-1 blockades were comparatively tested for prevention and/or treatment of PF induced by intratracheal BLM in mice.

In the prevention model, GF9 and GA31-LPC both effectively prevented PF at Day 22 (**Figure 4B**), while only GF9 but not GA31-LPC inhibited the earlier fibrotic process at Day 7 (not shown). In the treatment model, GF9 and GA31-LPC both effectively reversed PF. Interestingly, these data are in line with our recently published findings that TREM-1 blockade by either GF9 or GA31-LPC effectively prevented and reversed skin fibrosis induced by subcutaneous BLM in mice (adapted in **Figure 4C** for comparison purposes) and mitigated constitutive collagen synthesis and myofibroblast features in systemic sclerosis fibroblasts *in vitro* (44).

Thus, comparison of GF9 and GA31-LPC did not reveal significant differences between pan-TREM-1 and macrophage-restricted TREM-1 blockades for the prevention or treatment of PF. These findings are consistent with our previous data in experimental skin fibrosis (44) and RA (24–25), and are likely explained by the chronic nature of these inflammatory disorders.

## 4. Discussion

Myeloid cells, including neutrophils, monocytes and macrophages, are key players in the innate immunity response but may play dual roles in inflammation and diseases depending on the stage and context of the disease process (53–54). Complementing each other and closely interacting with each other, these cells efficiently defend the host against and eliminate pathogens (53). However, if not tightly regulated, the interactions between myeloid cells can be detrimental to the host by altering the balance between physiological and pathophysiological inflammatory responses (53). Myeloid cell function is closely controlled by TREM receptors including TREM-1 that amplifies the inflammation (2, 49).

Despite growing recognition of TREM-1 as an emerging target in cancer (3–4) and numerous other inflammation-associated diseases and disorders (reviewed in (5–9), the question whether pan-TREM-1 and cell-restricted TREM-1 (e.g., macrophage-restricted) blockades can differ in their therapeutic potential has never been addressed mainly due to the lack of proper methodology. By combining the ligand-independent SCHOOL mechanisms of TREM-1 inhibition by using the peptide sequence GF9 (**Figure 1A**) (59, 62) with nature-inspired LPC-based approach for delivery of drugs and imaging agents to macrophages via SR-A-mediated endocytosis (36, 59, 62, 83–84), we developed a first-in-class macrophage-restricted TREM-1 inhibitor GA31-LPC (**Figure 1B**). It should be noted that while in contrast to tissue-resident macrophages, neutrophils and monocytes do not express SR-A, the expression of SR-A is induced during differentiation of monocytes to macrophages (85). This suggests that due to the SR-A-mediated mechanism of delivery (36), GA31-LPC will inhibit TREM-1 on both tissue-resident and monocyte-derived types of macrophages while free peptide GF9 is not cell-specific and will inhibit TREM-1 on any of TREM-1-expressing cells (**Figure 1B**).

Here, I used GF9 and GA31-LPC to comparatively evaluate their therapeutic efficacy in animal models of cancer, sepsis, pulmonary inflammation and fibrosis. To my best knowledge, this is the first study to provide striking evidence for the previously unrecognized and seemingly paradoxical differences in the therapeutic activity and efficacy that pan-TREM-1 and macrophage-restricted TREM-1 blockades may exhibit in TREM-1-mediated inflammatory pathologies, depending on the disease, treatment timing and the type and stage of inflammatory response.

In cancer, pharmacological blockade of TREM-1 is a promising approach to target tumor-associated macrophages (TAM) (86–87), the major innate immune cells that infiltrate solid tumors and can form up to 50% of the tumor mass (86). TAMs play a pivotal role in tumor-associated chronic inflammation that promotes tumor progression (88). The clinical significance of TAMs is evidenced by the strong link between TAM number and density and a poor prognosis in 80% of the published cancer studies (89–92). On the other hand, cancer therapy (e.g., surgery, radiation, chemotherapy) can induce a strong acute inflammatory response by causing massive necrotic death of cancer cells and surrounding tissues and triggering a physiological inflammatory reaction similar to a wound-healing response (69, 93). If not resolved, therapy-induced acute inflammation may result in the failure of therapy, cancer recurrence and metastasis (73, 94). However, despite the well-established relationship between inflammation and cancer as well as encouraging data from multiple preclinical and observational population studies, anti-inflammatory therapeutics have failed to demonstrate significant benefit in the treatment or preventing recurrence of cancer (95–96). One example is canakinumab, a mAb that blocks the proinflammatory cytokine interleukin-1β, that failed in three Phase III trials for non-small cell lung cancer (97–98). Collectively, this highlights our poor understanding of the precise mechanisms and nuances involved in the complex relationship between inflammation and cancer, especially regarding different types and stages of inflammation as well as treatment types and timing. One of the key elements here is the balance between inflammation and resolution of inflammation, which if dysregulated, leads to disease pathology that may be associated with maladaptive immunity as a result of unresolved acute inflammation (99). This suggests pro-resolution strategies as a new therapeutic strategy that can be intrinsically more beneficial compared to conventional anti-inflammatory approaches.

In this study, in cancer mice with intact innate immunity but lacking functional adaptive immunity, a macrophage-restricted TREM-1 inhibitor GA31-LPC demonstrates remarkable anti-tumor activity when given after but not with chemotherapy (**Figure 2A**), suggesting that this activity is likely associated with resolution of acute innate inflammation caused by chemotherapy. Ongoing studies are focused on the determination of the therapeutic window for resolving cancer therapy-induced inflammation by macrophage-restricted TREM-1 blockade. In contrast, pan-TREM-1 inhibitor GF9 showed significant anti-tumor activity when given with but not after chemotherapy (**Figure 2A**), likely suggesting anti-inflammatory rather than pro-resolving mechanisms of its anti-tumor effect. Synergy of macrophage-restricted, but not pan-TREM-1 blockade with immunotherapy in fully immunocompetent mice (**Figure 2B**) adds further evidence to support the conclusion on the various contributions of myeloid cells to cancer progression (100).

While initially originated from studies on cancer mice treated with chemotherapy, the hypothesis that macrophage-restricted but not pan-TREM-1 blockade contributes to the resolution of acute inflammation has been further supported by key findings in other animal models of acute inflammatory response (sepsis and ALI). In these models, GA31-LPC demonstrated the striking and seemingly paradoxical dependence of its therapeutic efficacy on the timing of treatment initiation relative to challenge – the lower or no benefit when administered preventatively as compared to therapeutic treatment (**Figures 3 and 4A**). Importantly, the striking difference in the ability of GA31-LPC to protect septic mice from death (**Figure 3**) or inhibit infiltration of neutrophils into the injured lungs of LPS-challenged rats (**Figure 4A**) in preventive (1 h prior challenge) and therapeutic modes (1 h post challenge) is not likely to be explained by the half life of GA31-LPC in circulation since GA31-LPC exhibits biphasic elimination with a rapid phase for 1 h and a slow phase for 40 h (not shown).

Interestingly, GF9 and GA31-LPC were both effective in preventing and treating BLM-induced pulmonary fibrosis (**Figure 4B**) suggesting that in chronic inflammatory diseases, anti-inflammatory mechanisms of both pan-TREM-1 and macrophage-restricted TREM-1 blockades may contribute to these effects. This is in line with the findings in this study that GF9 and GA31-LPC alone show significant anti-tumor efficacy in fully immunocompetent mice (**Figure 2 B**). The data of our previous studies that GF9 and GA31-LPC effectively prevent and treat other chronic inflammatory diseases such as BLM-induced skin fibrosis (**Figure 4C**; adapted from (44)) and collagen-induced arthritis (24–25) further support this conclusion.

In summary, this proof-of-concept study is the first to demonstrate the potential use of macrophage-restricted TREM-1 inhibitory strategy for the fine-tuning of the innate immune response by targeting resolution rather than suppression of physiological inflammation in a variety of inflammatory diseases. In cancer, a macrophage-restricted TREM-1 inhibitor could be used in at least, three ways: (a) alone to suppress TAM-mediated chronic inflammation and inhibit cancer progression; (b) in combination with ICB to overcome resistance of hard-to-treat cancers such as pancreatic cancer and synergize with otherwise ineffective immunotherapy, and (c) to complement existing cancer therapies such as surgery, radiation, and chemotherapy and timely resolve the physiological acute inflammatory response caused by these therapies. These approaches can be deployed to prevent cancer recurrence and metastasis, increase the complete response rate, decrease the number of treatment cycles while lengthening intervals between treatments to improve quality of life, and extend overall survival.

## 5. Conclusions

To the best of my knowledge, this is the first time that the efficacy of pharmacological blockade of TREM-1 in animal models of TREM-1-mediated inflammation-associated diseases has been demonstrated to depend firstly, on the type of TREM-1 blocker used (pan-TREM-1 or macrophage-restricted TREM-1 inhibitor) and secondly, on the disease, the animal model studied, treatment timing and the type and stage of inflammatory response. As TREM-1 has been implicated as an inflammation amplifier involved in virtually all inflammatory conditions, the findings reported here offer novel insights into the intricate and interconnected nature of the physiological and pathophysiological inflammatory immune responses and open new avenues in further research on spatially and temporally distinct roles of myeloid cells in these processes. Further, these findings can be used not only to correctly interpret preclinical and clinical data on TREM-1, but also to design future synergistic therapeutic approaches to overcome the challenges of targeting inflammation in the treatment of cancer and other inflammatory diseases and disorders.

## Data availability statement

The original contributions presented in the study are included in the article. Further inquiries can be directed to the author.

## Ethics statement

The animal studies were performed in strict accordance with the recommendations in the Guide for the Care and Use of Laboratory Animals of the National Institutes of Health (NIH) and in the United States Department of Agriculture (USDA) Animal Welfare Act (9 CFR, Parts 1, 2, and 3). All experimental procedures were reviewed and approved by the Institutional Animal Care and Use Committees of the organizations listed for compliance with regulations prior to study initiation and all experiments were performed in accordance with the approved protocols.

## Author contributions

ABS: Conceptualization and study designs, Funding acquisition, Investigation, Methodology, Preparation and characterization of TREM-1 inhibitors, Analysis and interpretation of data, Validation, Visualization, Writing – original draft, review & editing.

## Funding

This work was supported in part by grants R44CA217400 from the National Cancer Institute of (NCI), R43AR078110 from the National Institute of Arthritis and Musculoskeletal and Skin Diseases (NIAMS), R43GM128369 from the National Institute of General Medical Sciences (NIGMS), R43HL165734 from the National Heart, Lung, and Blood Institute (NHLBI). NCI, NEI, NIGMS, and NHLBI are components of the National Institutes of Health (NIH).

## Acknowledgements

I would like to thank Dr. Daniel Von Hoff for his ongoing encouraging and inspiring support and for his invaluable advice on induction and maintenance therapies of pancreatic cancer. I am also grateful to Dr. Zu Shen for his critical reading of the manuscript and for interesting discussions.

## Funding

This work was supported in part by grants R44CA217400 from the National Cancer Institute of (NCI), R44EY034015 from the National Eye Institute (NEI), R43GM128369 from the National Institute of General Medical Sciences (NIGMS), R43HL165734 from the National Heart, Lung, and Blood Institute (NHLBI). NCI, NEI, NIGMS, and NHLBI are components of the National Institutes of Health (NIH).

## Conflict of Interest

The author Alexander B. Sigalov is employed by SignaBlok, Inc.

